# HyperMap: An Efficient Framework for Transferring Perturbation Responses Across Diverse Biological Contexts

**DOI:** 10.64898/2026.04.23.720505

**Authors:** Bhavya Dhaka, Jiahao Gao, Trey Ideker

## Abstract

Recent perturbation atlases profile transcriptional responses to thousands of targeted perturbations in a reference cell type. Generalising these datasets across lineages and individuals has been problematic, however, as similar baseline transcriptomes can yield highly divergent responses. To address this challenge, we present HyperMap, a meta-learning framework that translates existing atlases to predict perturbation responses in new biological contexts using a small number of perturbation “seeds.” Applied to CRISPR gene knockdowns in induced pluripotent stem cells, HyperMap accurately captures responses of new iPSC donors. It generalises to additional cell lines, perturbations by small-molecule drugs, and knockdowns not yet performed in any context. HyperMap is highly efficient, obtaining best-in-class predictions with one-eighth the parameters of typical foundation models. Integrating across atlases yields HyperMapDB, a complete 18×19,036 (cell-line × perturbation) matrix expanding current data by 27-fold. HyperMap enables predictive maps spanning the combinatorial space of biological contexts, gene knockdowns and drugs.

## Introduction

Measuring how genetic and chemical perturbations alter gene expression has long been a primary mode of understanding mechanisms of gene and drug action and their regulation^1–3^. Genetic perturbation expression profiling was greatly accelerated by the pooled approach known as “perturb-seq”^4^, which combines single-cell RNA sequencing with CRISPR (Clustered Regularly Interspaced Short Palindromic Repeats) to simultaneously profile the effects of many single-gene knockouts on genome-wide transcription patterns. Similarly, drug response profiling has expanded from bulk phenotypic measurements^5^ to single-cell transcriptomic readouts^6^, enabling detection of subtle and heterogeneous shifts in cell states. The growing throughput and cost-effectiveness of these methods have made it feasible for individual perturbation atlases to cover tens of thousands of perturbations^7,8^.

Computational modelling of perturbation responses has been pursued in at least two primary directions. The first attempts to use a limited set of experimentally profiled perturbations to predict responses to unseen perturbations. This aim has thus far been addressed by models that incorporate prior information from gene-gene functional interaction graphs^9^, gene-function text embeddings^10^, drug molecular structures^11^, or other biologically informed features^12^. As these sources of prior information are general, not tuned to a specific context, these methods generally do not capture context-specific transcriptional responses. The second direction attempts to predict cell states across diverse biological contexts. One set of methods uses generative modelling based on variational autoencoders (VAEs), which represent perturbed cell states as a function of baseline expression, perturbation inputs, and optional contextual covariates (e.g., cell type)^13^. Here, a challenge is that the Gaussian priors imposed by VAEs can constrain expressive power and lead to over-smoothed predicted responses^14^. A complementary approach is single-cell foundation modelling, in which training is performed on millions of cells across diverse tissues and perturbation conditions^15,16^. However, several studies have reported that such enormous models do not yet consistently outperform simpler methods^17,18^.

These challenges highlight the need for perturbation-response models that can quickly adapt to previously unseen cellular contexts without extensive new training. In this regard, meta-learning has emerged as a promising strategy for obtaining transferable representations in scenarios where data are scarce or costly to obtain^19^. Commonly described as “learning to learn,” meta-learning leverages experience across related tasks to rapidly adapt to new ones using only a few task-specific examples^20–22^. Meta-learning has shown strong success in computer vision and natural language processing and, more recently, in computational biology for applications including survival modelling, molecular property prediction, and drug response inference^23–27^. Related to meta-learning, the “hypernetwork” architecture enables context-dependent parameterisation by generating the weights of a primary neural network model using a second model conditioned on variable contexts^28^. This design allows regulatory differences between cell types to be captured directly in the model parameters, rather than forcing all contexts into a single fixed representation. Incorporating hypernetworks into meta-learning offers a principled way to achieve fast and flexible model adaptation across contexts.

Here we present HyperMap, a deep learning framework that predicts how genetic or chemical perturbations alter transcriptional states across cell types (Fig. 1a). Using meta-learning and hypernetworks, HyperMap trains on a reference perturbation atlas to learn shared response patterns, then dynamically adapts to new cellular contexts using a small set of observed perturbations as seeds. These seed perturbations provide sufficient information for the model to predict responses for most remaining perturbations in that context with high performance. Unlike methods requiring fixed gene interaction graphs or extensive priors, HyperMap learns directly from data and can make accurate predictions even when experimental coverage is sparse. This behaviour enables perturbation prediction across diverse cell types and previously unobserved biological conditions, with applications spanning from basic cell biology to translational medicine.

**Fig. 1.**
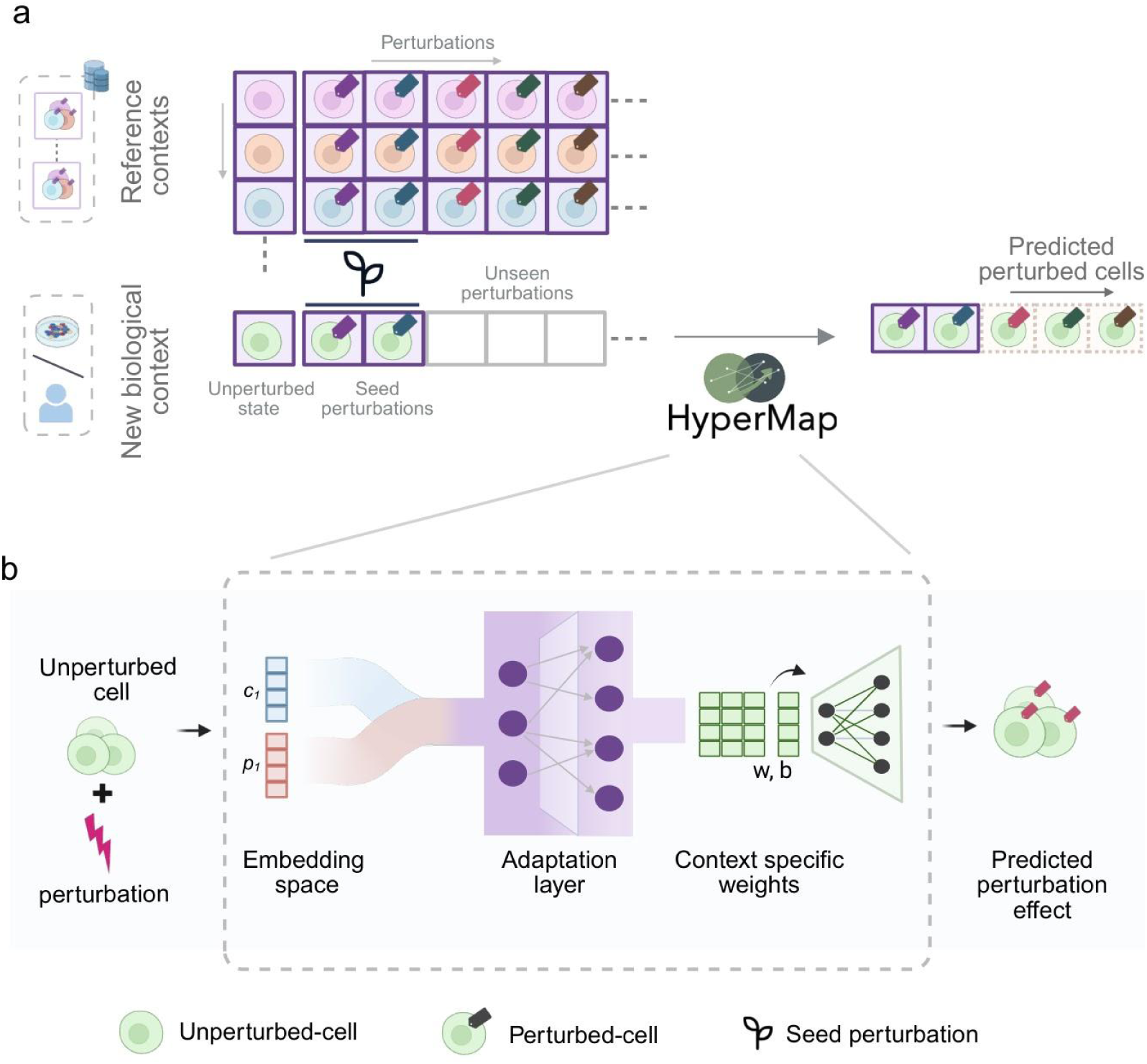
HyperMap overview and model architecture. **a)** Schematic illustrating how HyperMap transfers perturbation states from previously measured reference contexts (top rows) to a new biological context (bottom row). Reference contexts consist of perturbation-response profiles, e.g. from large public perturb-seq studies spanning either genetic or chemical perturbations (columns). The new context contains only a small number of shared seed perturbations, which HyperMap uses to adapt information learned from the reference contexts and generate predicted responses for unseen perturbations. **b)** Diagram of the HyperMap architecture. An unperturbed cell and a perturbation identity are encoded into a shared embedding space, forming the unique context for the adaptation layer. This layer generates context-specific weights and biases (w, b) that parameterise the downstream predictor, which outputs the predicted post-perturbation effects for that context.

## Results

### A context-aware meta-learning model of transcriptional perturbation

HyperMap transfers a model of cell transcriptional state to new contexts through its context-dependent architecture (Fig 1b, Methods). A perturbation response is predicted from two types of information, reflecting (1) the cell-type context and (2) the applied perturbation. The cell type is represented by its unperturbed gene-expression profile, encoding the baseline regulatory state, whereas the perturbation is represented as either a one-hot vector or a learned gene-based embedding from GenePT^29^, which embeds each gene using its NCBI descriptive summary text to capture functional features. The cell type and perturbation embeddings are combined to create a unique context representation specific to that cell-perturbation pair.

Next, this context representation is passed to the adaptation layer, the central component of HyperMap (Fig 1b). The adaptation layer contains a hypernetwork^28^ that takes the cell-perturbation embedding and generates a set of weights and biases for the downstream perturbation-response predictor. When applied to a new cellular context, a small set of seed perturbations fine-tunes the adaptation layer to align with that context’s response patterns. Through this mechanism, HyperMap can adjust to a new cellular context without retraining, since its parameters are generated dynamically from the input rather than being fixed, allowing it to capture transcriptional responses to the same perturbation across different contexts. Rapid adaptation is further enabled by the meta-learning strategy used during training (Extended Fig. 1a,b).

### Control-state similarity does not predict perturbation response

A common assumption in perturbation biology is that cell types with similar baseline states will exhibit similar responses to perturbation. To assess this assumption, we analysed a large-scale induced pluripotent stem cell (iPSC) perturb-seq atlas consisting of 436 CRISPR gene knockdown perturbations profiled across 10 donors^30^. For each donor pair, we computed baseline transcriptomic similarity and perturbation response similarity under each CRISPR perturbation (Methods), finding no association between the two (Fig. 2a).

**Fig. 2.**
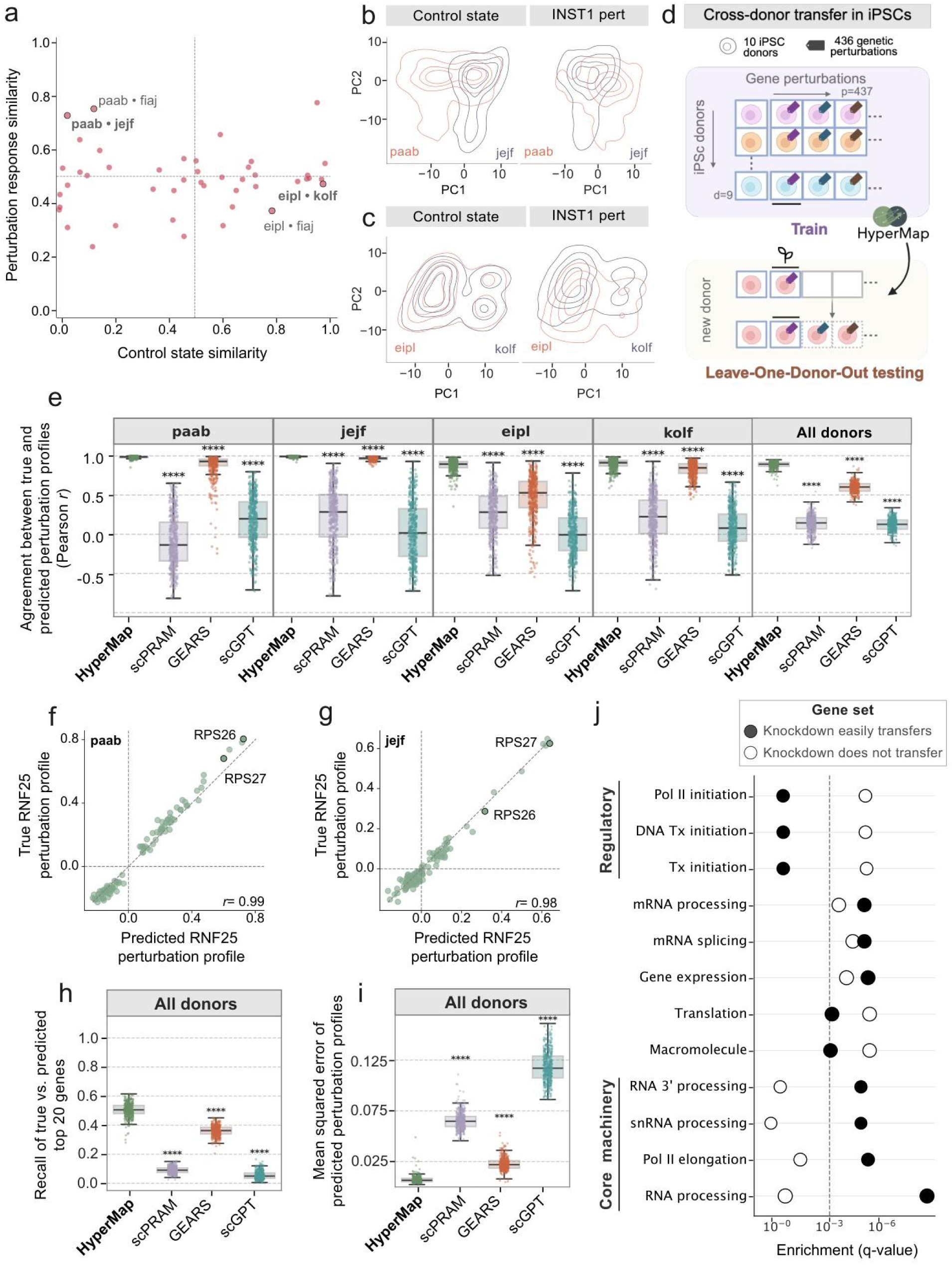
Cross-donor transfer of molecular states in human iPSCs. **a)** Scatter plot showing donor-pair (points) similarity in the control state (x-axis) versus similarity in the perturbation response (y-axis) across 436 perturbations. Dotted horizontal and vertical lines indicate the similarity midpoint of 0.5. **b,c)** Contour plots for donor pairs, paab-jejf (b) and eipl-kolf (c), showing the distributions of single-cell transcriptional states for each donor (top two principal components, PCs) under the control state (left) or the INST1 perturbation (right). **d)** Approach used for HyperMap training and testing. HyperMap is trained on all donors except one, and predictions are generated for the held-out donor. For the held-out donor, the model is adapted using 10 seed perturbations and evaluated on the remaining 426 perturbations. **e)** Model performance across perturbations for individual donors. The y-axis shows the Pearson correlation between the predicted and true change in expression, computed for the top 20 differentially expressed genes. The x-axis lists the models used to generate predictions. Each point represents one perturbation (n = 426). The boxes represent the interquartile range (IQR) bisected by the median, and whiskers represent the maximum and minimum range of the data that do not exceed 1.5 times the IQR. Statistical significance was assessed using a two-sided Mann-Whitney U test (**** p < 0.0001). **f-g)** Correlation between predicted and true RNF25 perturbation profiles for two example donors (paab and jejf). Each point represents an individual gene (the top 100 response genes are shown). The dashed line indicates the identity line; Pearson correlation coefficients are shown. **h)** Recovery of top perturbation-responsive genes across all donors. The y-axis shows the fraction of correctly recovered top-ranked perturbation-responsive genes relative to observed top-ranked genes, and the x-axis lists the models used to generate predictions. Each point represents one perturbation, averaged across donors. **i)** Error of predicted perturbation profiles across all donors. The y-axis shows the mean squared error between predicted and true perturbation profiles. Other details are as in panel h. **j)** Major functions of gene knockdown, grouped by transferability of their perturbation response profiles. Filled circles (Knockdown easily transfers) indicate gene sets whose knockdown responses are consistently well predicted across donors (top 25% by transferability); open circles (Knockdown does not transfer) indicate gene sets that are poorly predicted across donors (bottom 25% by transferability). Each circle represents one gene set; rows show Gene Ontology Biological Process terms enriched in one or both groups. The dashed vertical line represents the GO term enrichment significance threshold (q-value = 0.001). Abbreviations: Tx, transcription. Full names for each term are documented in Extended Table 1.

This disconnect is illustrated by perturbation of INTS1, a core component of the Integrator complex. Donors *paab* and *jejf* have low baseline transcriptional similarity yet show converged responses to INTS1 perturbation (Fig. 2b), whereas donors *eipl* and *kolf* have high baseline similarity yet show diverged responses (Fig. 2c). As a necessary control, we found that iPSC lines derived from the same donor show high similarity in both control state and perturbation responses (Extended Fig. 2a,b). These observations highlight that relying on the “closest” baseline neighbour does not reliably predict perturbation responses, underscoring the need for models that explicitly learn context-specific regulatory structures.

### HyperMap transfers perturbation responses across diverse iPSC donors

We next tested HyperMap’s ability to capture donor-specific perturbation effects. For this, we trained the model using data from nine donors, then adapted it to the held-out donor using ten seed perturbations (Fig. 2d). The model was then used to predict that donor’s transcriptional responses to each of the remaining 426 perturbations (Fig. 2e, Methods). In evaluating model performance, we found that the correlation between predicted and true expression profiles ranged from 0.72 to 0.99 across individual donors (Fig. 2e, Extended Fig 2). For example, in perturbing RNF25, an E3 ubiquitin ligase, HyperMap captured both the direction and magnitude of gene-expression changes, including concordant upregulation of the ribosomal proteins RPS26 and RPS27 (Fig. 2f,g).

This performance was compared to a panel of state-of-the-art perturbation response prediction models (Methods), including scPRAM^31^, which leverages variational autoencoders and optimal transport; GEARS^9^, a graph-based method using Gene Ontology^32,33^ to predict perturbation responses; and scGPT^15^, a foundation model pretrained on transcriptomes of millions of cells. HyperMap achieved the highest predictive performance among these models (Fig. 2e), and more closely matched observed perturbation responses than either the mean perturbation effect across training donors or the average unperturbed transcriptional state (Extended Fig. 2d-g), confirming that its predictions reflect genuine context-specific perturbation effects rather than generic or background signals.

We also evaluated HyperMap using performance metrics other than Pearson correlation, such as the recovery of the top perturbation-responsive genes. HyperMap achieved the highest recovery (Fig. 2h) with the lowest mean squared prediction error (Fig. 2i) across all models. GEARS, which explicitly incorporates gene-gene interaction graphs, showed the second-best recovery, whereas foundation models such as scGPT exhibited a marked reduction in recovery. This pattern suggests that explicitly modelling local gene relationships can aid in the identification of perturbation-specific targets compared to approaches that rely primarily on global expression representations.

To investigate whether particular gene perturbations generalise better across donors, we ranked gene knockdowns by their accuracy of HyperMap prediction, defining the top versus bottom 25% as strongly versus weakly transferable, respectively. Strongly transferable perturbations were enriched for core cellular machinery, including RNA processing and elongation, whereas weakly transferable targets were enriched in regulatory functions such as transcription initiation, which are more context-dependent^34^ (Fig. 2j, Extended Table 1). This pattern was not explained by baseline expression levels, as strongly transferable genes were not more highly expressed (Extended Fig 2h). These results suggest that the ease of perturbation transfer reflects biological variability in regulatory control rather than a technical artefact.

### Transfer of transcriptional response extends to diverse cell lines and perturbation types

We next sought to assess HyperMap in an expanded array of biological contexts, moving from iPSC donors to panels of distinct human cell lines and from CRISPR gene knockdowns to small-molecule drug treatments. First, we accessed the SciPlex3 dataset^6^, comprising 188 compounds profiled across A549, K562, and MCF-7 tumour cell lines (Fig. 3a). HyperMap was trained on two cell lines and evaluated on the held-out third, where 20 seed perturbations were used to fine-tune the model, leaving 168 compounds for prediction. Given equivalent performance with one-hot encodings in the genetic setting (Extended Fig. 3a-c), we adopted these for chemical perturbations. HyperMap achieved a median Pearson correlation of approximately 0.46, which was significantly higher than scPRAM (Fig. 3b), accompanied by higher recovery of top perturbation response genes (Fig. 3c) and lower mean squared error (Fig. 3d). GEARS and scGPT, which require gene-centric perturbation representations, could not be applied in this setting. HyperMap achieved strong predictive performance across all cell lines, as illustrated by streptozotocin responses in A549 (r= 0.84; Fig. 3e), K562 (r= 0.50; Fig. 3f), and MCF-7 (r= 0.62; Fig. 3g). The lower accuracy in K562 likely reflects its distinct hematopoietic lineage relative to the epithelial training lines, suggesting substantial differences in underlying regulatory programs.

**Fig. 3.**
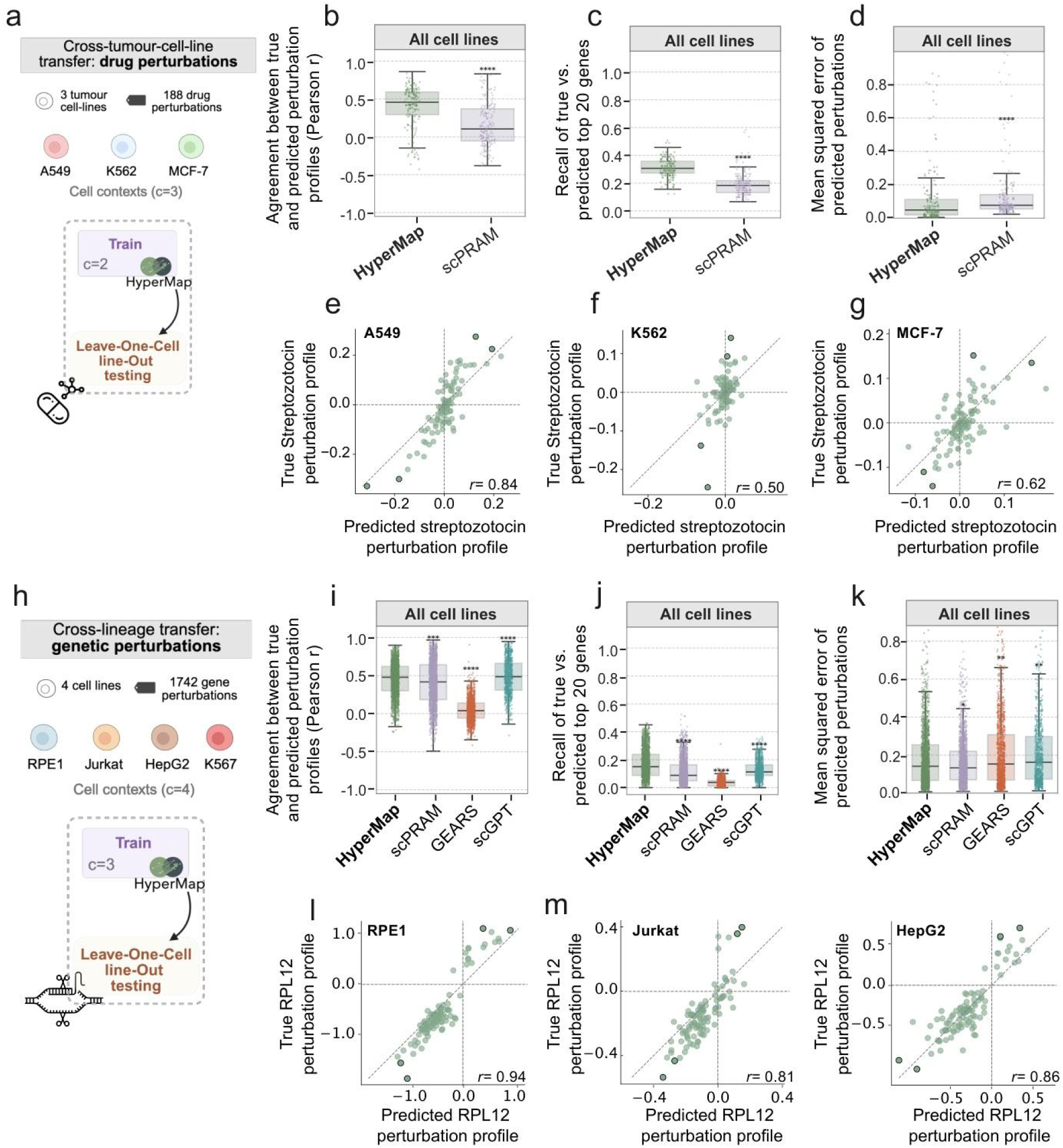
Performance of HyperMap for transfer of genetic and chemical perturbations across tumour cell lines. **a)** Approach used for HyperMap training and testing for cross-cell-line transfer of drug perturbations (n=188). HyperMap is trained on all cell lines except one, and predictions are generated for the held-out cell line (A549, K562, or MCF-7). For the held-out cell line, the model is adapted using 20 seed perturbations and evaluated on the remaining 168 drug perturbations. **b-d)** Model performance aggregated across all cell lines for drug perturbations. Each point represents one drug perturbation (n = 168). Statistical significance was assessed using a two-sided Mann-Whitney U test (* p < 0.05; ** p < 0.01; *** p < 0.001; **** p < 0.0001). Other details are as in Fig. 2e. b) Pearson correlation between the predicted and true change in expression, computed for the top 20 differentially expressed genes. c) Fraction of correctly recovered top response genes relative to observed. d) Error of predicted perturbation profiles. **e-g)** Correlation between predicted and true perturbation profiles for the streptozotocin drug perturbation in three cell lines (e A549, f K562, g MCF-7). Each point represents an individual gene (the top 100 genes are shown). The dashed line indicates the identity line; Pearson correlation coefficients are shown. **h)** Approach used for HyperMap training and testing for cross-cell-line transfer of genetic perturbations (n = 1,742). HyperMap is trained on all cell lines except one, and predictions are generated for the held-out cell line (RPE1, Jurkat, K562, or HepG2). For the held-out cell line, the model is adapted using 20 seed perturbations and evaluated on the remaining 1,722 gene perturbations. **i-k)** Model performance aggregated across all cell lines for genetic perturbations. Each point represents one gene perturbation (n = 1,722). Other details are as in panel b-d. **l-n)** Correlation between predicted and true perturbation profiles for the RPL12 genetic perturbation in three example cell lines (l RPE1, m Jurkat, n K562). Each point represents an individual gene (the top 100 genes are shown). The dashed line indicates the identity line; Pearson correlation coefficients are shown.

We next analyzed two complementary perturb-seq resources^7,35^, spanning CRISPR knockdowns of 1,742 essential genes in epithelial (RPE1), hepatic (HepG2), hematopoietic (K562), or lymphoid (Jurkat) lineages (Fig. 3h). HyperMap was trained on three of these cell lines and evaluated on the held-out fourth, where 20 seed perturbations were used to fine-tune the model, leaving 1,722 gene knockdown responses for prediction. Despite pronounced lineage and morphological differences across cell lines, HyperMap had significant ability to predict responses to perturbation (r= 0.47, p= 0.03), achieving performance comparable to or better than GEARS, scPRAM, and scGPT across all metrics (Fig. 3i), including recovery of top response genes (Fig. 3j) and prediction error (Fig. 3k). This strong predictive performance was illustrated by knockdown of RPL12, a core ribosomal protein, in RPE1 (r= 0.94; Fig. 3l), Jurkat (r= 0.81; Fig. 3m), HepG2 (r= 0.86; Fig. 3n), and K562 (r= 0.65; Extended Fig. 3d).

### HyperMap extends to perturbations not yet performed in any context

Beyond cross-context transfer, a complementary challenge is to predict responses for new perturbations not yet observed during training. To evaluate this capability, we applied HyperMap to the iPSC perturb-seq dataset in a modified leave-one-donor-out design (Fig. 4a). HyperMap was trained on nine donors using a restricted set of 200 gene knockdown perturbations, adapted to a held-out donor using seed perturbations, then used to predict responses for 236 perturbations withheld during training and an additional 18,609 protein-coding perturbations not yet assayed experimentally. GEARS and scGPT, which can also generate predictions for unseen perturbations, were evaluated on the same 236 withheld perturbations under identical training restrictions; scPRAM requires reference empirical perturbation data, thus could not be applied. HyperMap achieved consistently high agreement between predicted and observed perturbation profiles, comparable to the full training set performance, and matching or exceeding GEARS and scGPT (Fig. 4b).

**Fig. 4.**
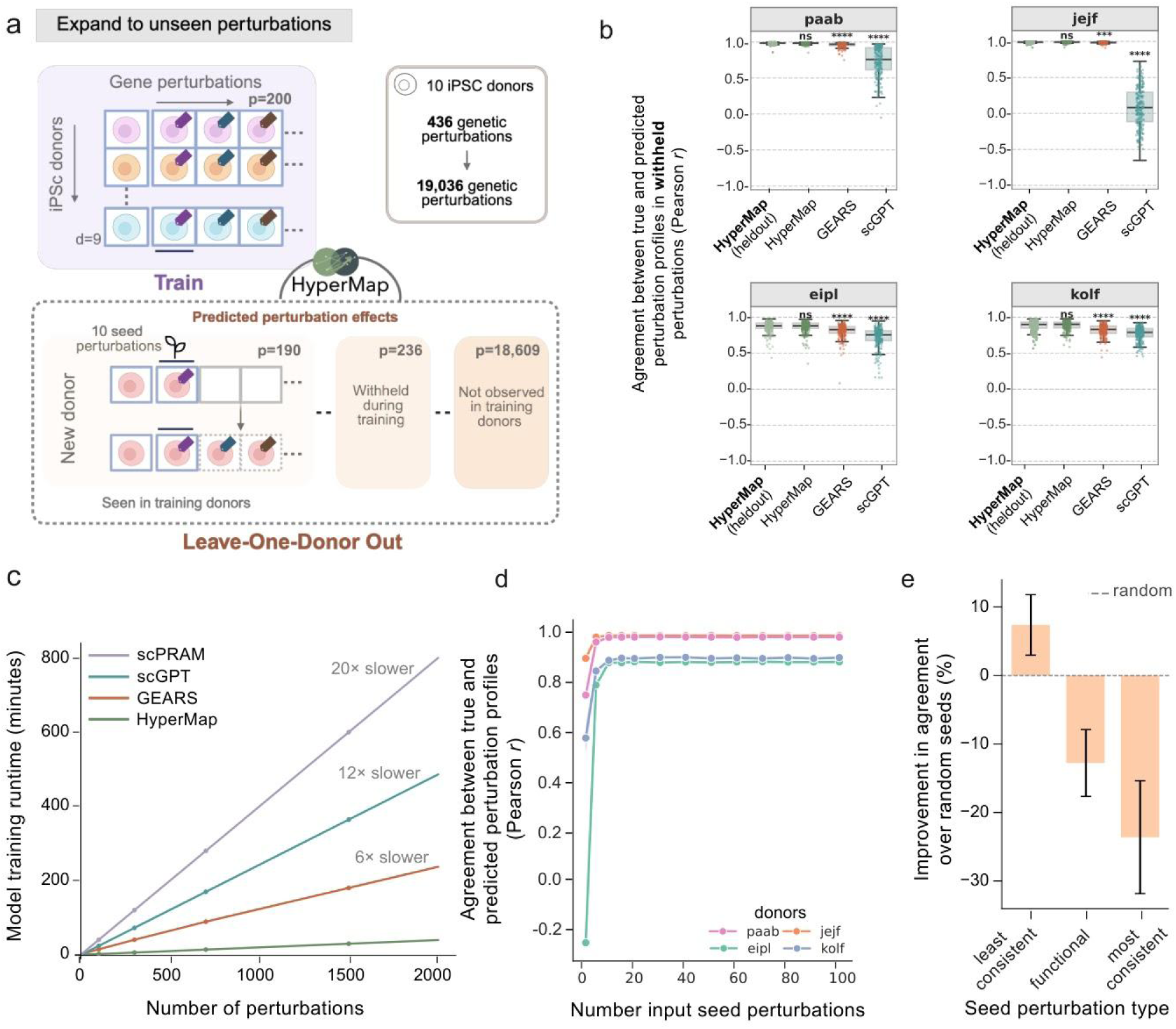
Predicting responses to unseen perturbations and analysis of scalability and seed efficiency. **a)** Schematic illustrating the HyperMap framework for predicting unseen genetic perturbations. Training uses 200 gene perturbations across nine donors; the held-out donor is adapted with 10 seed perturbations and evaluated on 236 withheld and 18,600 experimentally unobserved perturbations (19,036 total). **b)** Model performance across held-out perturbations (points, n=236) for four example donors. The y-axis shows the Pearson correlation between the predicted and true change in expression, computed for the top 20 differentially expressed genes. Statistical significance was assessed using a two-sided Mann-Whitney U test (**** p < 0.0001; ns non-significant). Other details are as in Fig. 2e. **c)** Model training runtime as a function of the number of perturbations. The x-axis shows the number of perturbations used for training, and the y-axis shows model training time in minutes. Each trace represents one model (scPRAM, scGPT, GEARS, HyperMap). Points indicate measured runtimes (100, 300, 700, 1500) and lines indicate linear interpolation between measurements. **d)** Model performance across perturbations for four example donors (colored traces) as a function of the number of input seed perturbations. The x-axis shows the number of seed perturbations used for adaptation, and the y-axis shows the Pearson correlation between the predicted and true changes in expression, computed for the top 20 differentially expressed genes. Results are averaged over 10 runs with randomly sampled seed perturbations; shaded regions indicate variance across runs. **e)** Improvement in model performance over random seeds, stratified by seed perturbation type. The x-axis lists seed perturbation types (least consistent, functional, most consistent), and the y-axis shows the percent improvement in prediction performance relative to randomly selected seeds (n=10). Bars indicate the mean improvement across donors, and error bars indicate variance across donors.

### HyperMap is computationally efficient and benefits from seed selection

We next evaluated the computational efficiency of HyperMap, including model training requirements and how predictive performance is influenced by the number and choice of seed perturbations. HyperMap training time scaled linearly with dataset size, requiring approximately 20 minutes per 1000 perturbations on a single GPU 32GB (Fig. 4c, Methods). This result was favourable in comparison to the panel of alternative models, which were between 6- and 20-fold slower (Fig. 4c). The relative efficiency of HyperMap relates to the meta-learning strategy and hypernetwork architecture, which enable powerful context adaptation while keeping the core neural network simple (Extended Fig. 3f-g).

To examine how adaptation performance depends on seed perturbations, we progressively increased the number of seeds used to adapt HyperMap to a held-out iPSC donor. Prediction accuracy increased rapidly and plateaued after approximately 20 perturbations (Fig. 4d), indicating that accurate transfer requires relatively limited adaptation data. Donors whose transcriptional profiles more closely resembled the training set (*paab* and *jejf*) reached predictive accuracy of r = 0.82 with just a single seed, suggesting that biological proximity to reference contexts further reduces data requirements. Seed choice also mattered: compared to randomly selected seed perturbations, selecting gene knockdowns with high response variability (low consistency) across training donors improved correlation by ∼8%, while selecting gene knockdowns related to core iPSC functionality or with high consistency reduced performance by ∼12% and ∼23%, respectively (Fig. 4e). Together, these results show that both the number and biological properties of seeds critically shape transfer performance.

### HyperMapDB: A complete resource of genome-wide knockdown responses for 18 contexts

Given HyperMap’s high efficiency and its demonstrated ability to transfer perturbation responses across biological contexts and to unseen perturbations, we sought to integrate and gap-fill across all available perturb-seq atlases to yield a unified, complete resource (Fig 5a). We curated perturb-seq datasets from independent screens^36,37^ yielding data from 18 contexts, including HepG2^35^, hESC^38^, Jurkat^35^, K562^7^, Luhmes^40^, MCF10A^41^, neurons^42^, RPE1^7^, and the iPSC donors^30^ (Extended Table 2). The number of perturbations ranged from 14 to 2,317 per context (Fig 5b), with a combined total of 2,531 distinct single gene knockdowns experimentally observed in at least one context and covering only 12,861 of the 45,558 possible context-by-perturbation combinations. We retrained HyperMap on the full collection, predicted all unobserved cell states within this set, and further expanded predictions to the complete protein-coding genome, yielding transcriptomic cell states for 19,036 perturbations across all 18 contexts (Fig. 5b) representing 342,648 context-by-perturbation combinations, a 27-fold increase over the existing atlases^7,39,43^ (Fig. 5a,b).

**Fig. 5.**
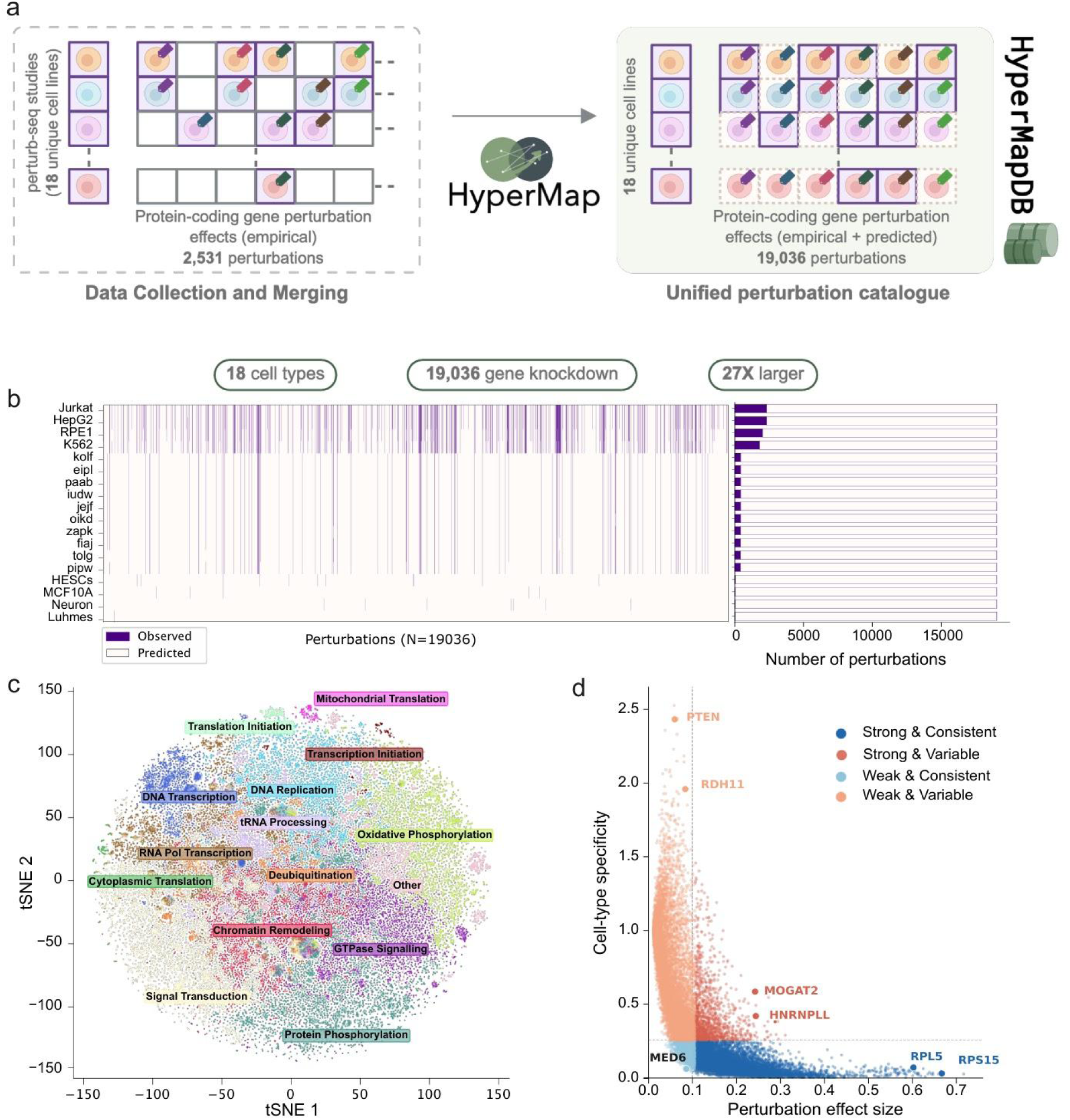
Predicting responses to unseen perturbations yields a complete context-by-perturbation atlas. **a)** Schematic illustrating the construction of the unified perturbation catalogue. Perturb-seq studies spanning 18 unique cell lines with empirically measured protein-coding gene perturbation effects are integrated and expanded using HyperMap to generate a unified catalogue of 19,036 perturbations. **b)** Coverage of observed (dark purple) and predicted (white) perturbations across 18 cell types in the unified catalogue. Each column represents one perturbation (N = 19,036), and each row represents one cell type. The bar plot on the right shows the total number of perturbations per cell type, stacked by observed and predicted. **c)** tSNE embedding of predicted perturbation response profiles across all cell types and perturbations in the unified catalogue. Each point represents one perturbation-cell type pair, colored by Leiden cluster. Labels indicate enriched biological processes within the perturbations of each cluster. **d)** Scatter plot showing perturbation effect size (x-axis) versus cell-type specificity (y-axis) for all perturbations in the unified catalogue. Each point is a perturbation, colored by perturbation class: Strong & Consistent (dark blue), Strong & Variable (dark orange), Weak & Consistent (light blue), and Weak & Variable (light orange). Representative gene knockdowns are labelled.

Projecting this response matrix into two dimensions (t-SNE, Methods) revealed that perturbation identity is the dominant organising principle, with perturbed genes grouping into coherent biological processes including DNA transcription, chromatin remodelling, and oxidative phosphorylation (Fig. 5c). For a minority of perturbations, cell-type identity overrode this pattern, forming isolated context-specific islands (Fig. 5c). Inspection of predicted response profiles revealed that neurons and Luhmes cells, derived from entirely independent datasets, displayed similar response patterns, consistent with their shared neuronal identity (Extended Fig. 4a). Among iPSC donors, two groups with broadly opposing response directions were apparent, recapitulating anti-correlated perturbation responses in the experimentally observed data (Extended Fig. 4b).

Decomposing perturbation magnitude and cell-type specificity across all 19,036 predicted perturbation responses revealed distinct modes of gene function (Fig. 5d). Core translational machinery, including RPS15 and RPL5, produced large, uniform effects across all cell types, reflecting their universal essentiality^44,45^. HNRNPLL displayed strong but highly context-restricted effects, in line with its known role in lymphocyte-specific splicing^46^. Similarly, PTEN and RDH11 showed highly context-dependent effects consistent with their tissue-specific activity^47,48^. MED6, a core component of the Mediator transcriptional complex, showed limited cell-type specificity^49^.

## Discussion

Accurately predicting cellular responses to perturbation is a central requirement for building predictive models of cell state and, ultimately, a virtual cell. Here we introduce HyperMap, a context-aware meta-learning framework that predicts transcriptional responses to genetic and chemical perturbations and rapidly adapts to new cellular contexts using only a small number of example perturbation “seeds”. A key observation motivating this approach is that baseline transcriptomic similarity between cellular contexts does not predict similarity in their perturbation responses (Fig. 2a,b). This finding highlights the need for models that learn context-specific regulatory structure rather than relying on proximity in expression space.

Previous approaches to cell state prediction such as foundation modelscan achieve strong performance but often require large training datasets, complex priors, or computationally intensive training procedures that limit scalability and deployment. In contrast, HyperMap uses a relatively simple core neural network and learns context-specific adaptation through a hypernetwork trained via meta-learning. Despite this minimal design, HyperMap matches or exceeds the performance of more complex models across multiple settings while scaling more favourably with dataset size.

Consistent with meta-learning theory, effective generalisation depends on exposure to diverse training tasks, each corresponding to a perturbation applied within a specific cellular background (Extended Fig 1). Sophisticated perturbation embeddings were not required for strong performance: replacing pretrained perturbation representations with simple one-hot encodings did not reduce accuracy (Extended Fig. 3). Together with the observation that seed perturbations producing diverse transcriptional responses improve model adaptation, these results indicate that both context-dependent learning and the biological diversity of seed perturbations are critical for effective transfer.

We also observed systematic differences in perturbation transferability across genes. Perturbations targeting core cellular machinery, such as RNA processing or transcription elongation, showed high transferability across contexts, whereas perturbations affecting regulatory or cell-type-specific processes were less predictable (Fig. 2j). This pattern highlights an inherent limitation of cross-context transfer: when regulatory programs differ substantially between reference and target contexts, predictive accuracy declines. As perturbation datasets expand to include additional tissues, cell types and disease contexts, transfer performance should improve naturally without requiring heavier modelling assumptions or curated priors.

Like many hypernetwork-based approaches, HyperMap currently offers limited interpretability at the level of learned parameters. More broadly, our study suggests that scalable models of cellular state and response may emerge not from increasingly complex architectures, but from frameworks that efficiently learn how regulatory programs adapt across biological contexts.

## Extended Data

**Extended Data Fig. 1.**
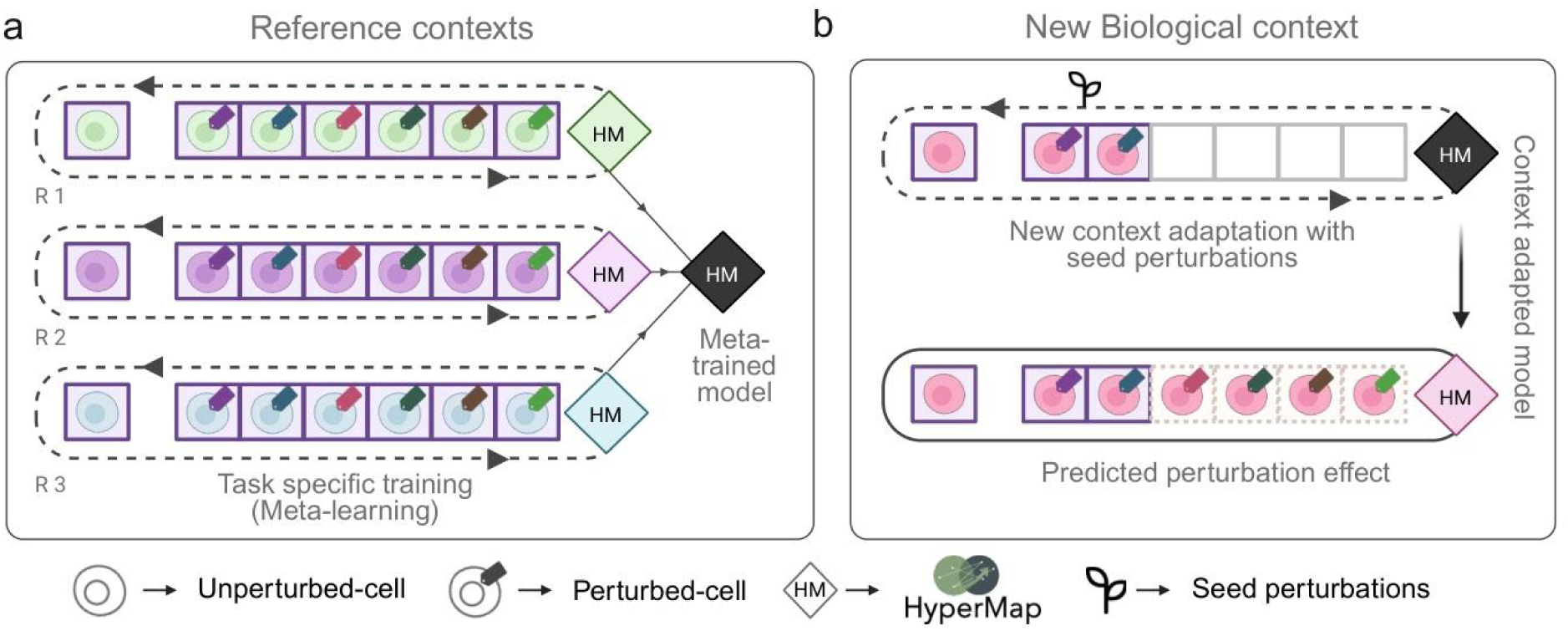
Meta-learning training strategy and new context adaptation in HyperMap. **a)** Schematic illustrating HyperMap training across three reference contexts (R1, R2, R3). Each reference context contains an unperturbed cell (leftmost) paired with a series of perturbed cells (columns), with a context-specific HyperMap model (colored diamond, HM) trained per context. A single meta-trained model (black diamond, HM) is produced by aggregating information across all reference contexts. **b)** Schematic illustrating adaptation of the meta-trained model to a new biological context. Seed perturbations adapt the meta-trained model to a context-adapted model (colored diamond, HM), which is then used to predict perturbation effects for unseen perturbations in the new context (dashed outline).

**Extended Data Fig. 2.**
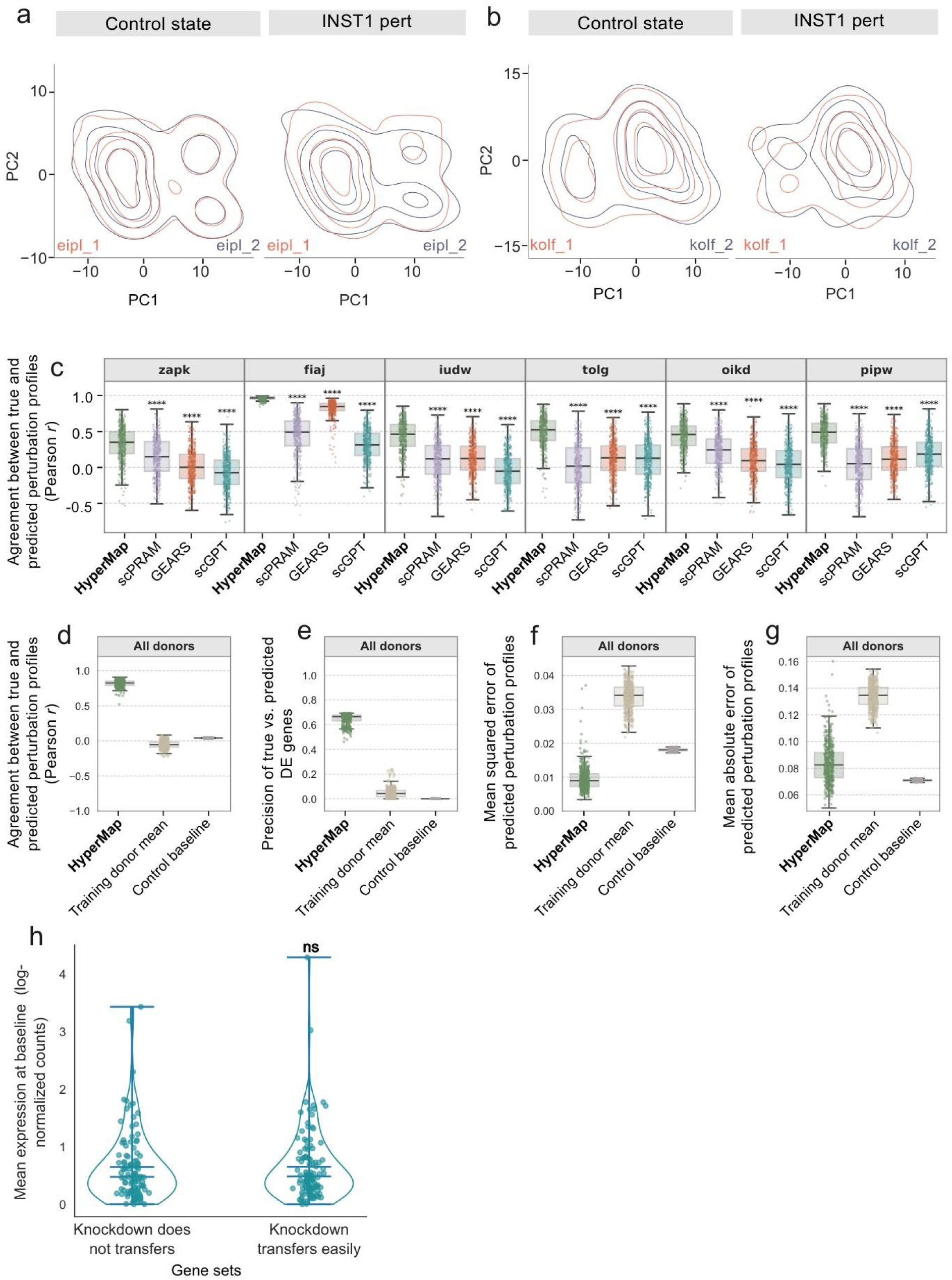
Cross-donor transfer of molecular states in human iPSCs a,b) Contour plots for two iPSC lines derived from the same donor, eipl_1 and eipl_2 **(a)** and kolf_1 and kolf_2 **(b)**, showing the distributions of single-cell transcriptional states for each line (top two principal components, PCs) under the control state (left) or the INST1 perturbation (right). **c)** Model performance across perturbations for six representative donors. The y-axis shows the Pearson correlation between the predicted and true change in expression, computed for the top 20 differentially expressed genes. The x-axis lists the models used to generate predictions. Each point represents one perturbation. The boxes represent the interquartile range (IQR) bisected by the median, and whiskers represent the maximum and minimum range of the data that do not exceed 1.5 times the IQR. Statistical significance was assessed using a two-sided Mann-Whitney U test (**** p < 0.0001). **d-g)** Model performance of HyperMap compared against two baselines (training donor mean and control baseline) aggregated across all donors. **d)** Pearson correlation between the predicted and true change in expression, computed for the top 20 differentially expressed genes. **e)** Precision in recovery of differentially expressed (DE) genes across all donors. The y-axis shows the fraction of correctly predicted DE genes relative to observed DE genes. **f)** Error of predicted perturbation profiles across all donors. The y-axis shows the mean squared error between predicted and true perturbation profiles. **g)** Mean absolute error of predicted perturbation profiles across all donors. Other details are as in panel d. **h)** Mean baseline expression of gene sets grouped by transferability of their perturbation response profiles. The x-axis shows gene set groups (Knockdown does not transfer; Knockdown transfers easily), and the y-axis shows mean expression at baseline (log-normalised counts). Each point represents one gene. Statistical significance was assessed using a two-sided Mann-Whitney U test (ns, not significant).

**Extended Data Fig. 3.**
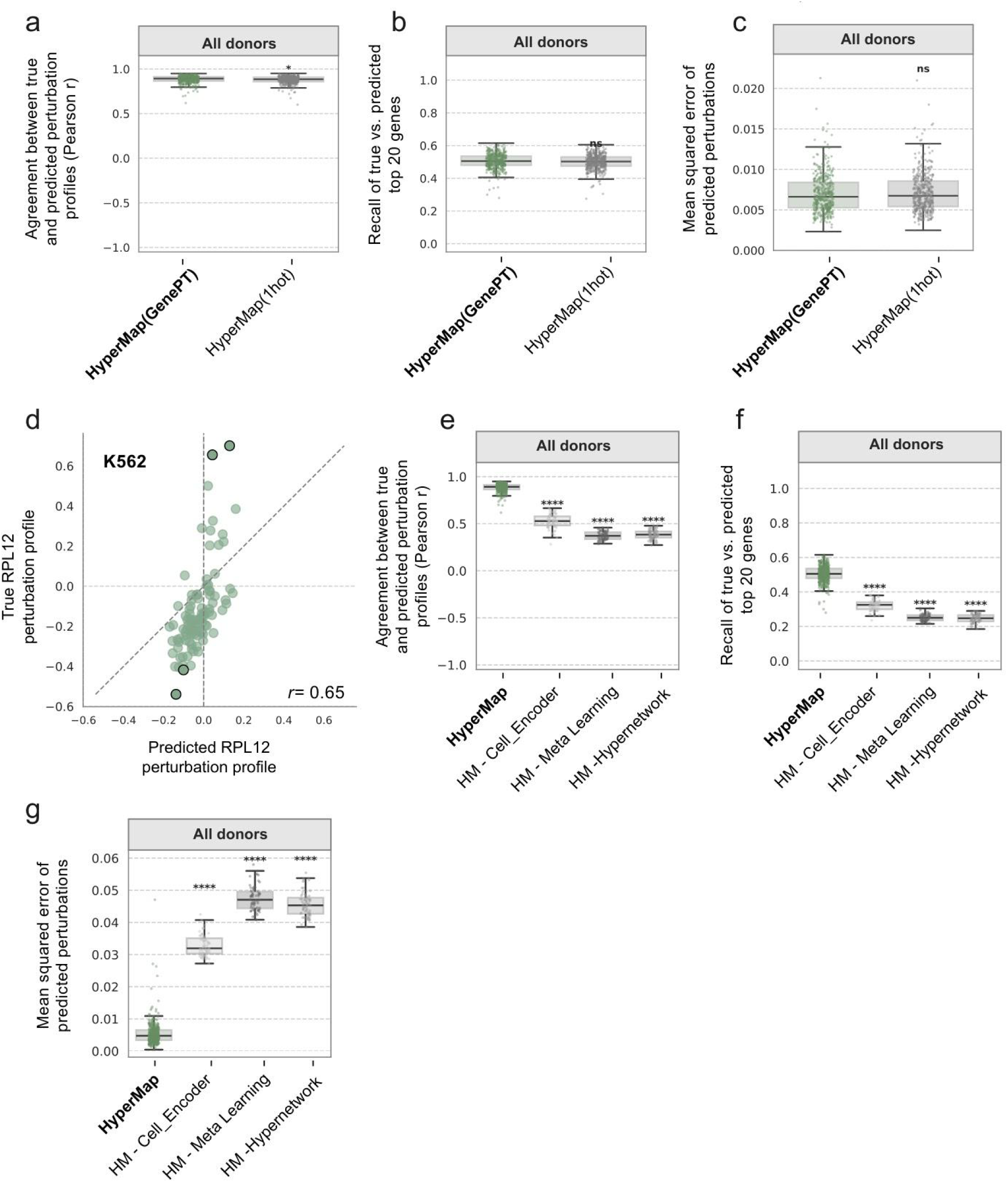
Perturbation encoding and ablation analysis of HyperMap components. a-c) Comparison of HyperMap performance using GenePT embeddings versus one-hot encodings for perturbation representation, aggregated across all donors. **a)** Pearson correlation between the predicted and true change in expression, computed for the top 20 differentially expressed genes. **b)** Recall of true versus predicted top 20 genes across all donors. **c)** Mean squared error of predicted perturbation profiles across all donors. Statistical significance was assessed using a two-sided Mann-Whitney U test (* p < 0.05; ns, not significant). Other details are as in Fig. 2d. **d)** Correlation between predicted and true RPL12 perturbation profiles for the K562 cell line. Each point represents an individual gene (the top 100 genes are shown). The dashed line indicates the identity line; the Pearson correlation coefficient is shown. **e-g)** Ablation analysis of HyperMap architectural components aggregated across all donors. Models compared are the full HyperMap and three ablated variants: HM - Cell_Encoder (without cell encoder), HM - Meta Learning (without meta-learning), and HM - Hypernetwork (without hypernetwork). **e)** Pearson correlation between the predicted and true change in expression, computed for the top 20 differentially expressed genes. **f)** Recall of true versus predicted top 20 genes across all donors. **g)** Mean squared error of predicted perturbation profiles across all donors. The boxes represent the interquartile range (IQR) bisected by the median, and whiskers represent the maximum and minimum range of the data that do not exceed 1.5 times the IQR. Statistical significance was assessed using a two-sided Mann-Whitney U test (**** p < 0.0001).

**Extended Data Fig. 4.**
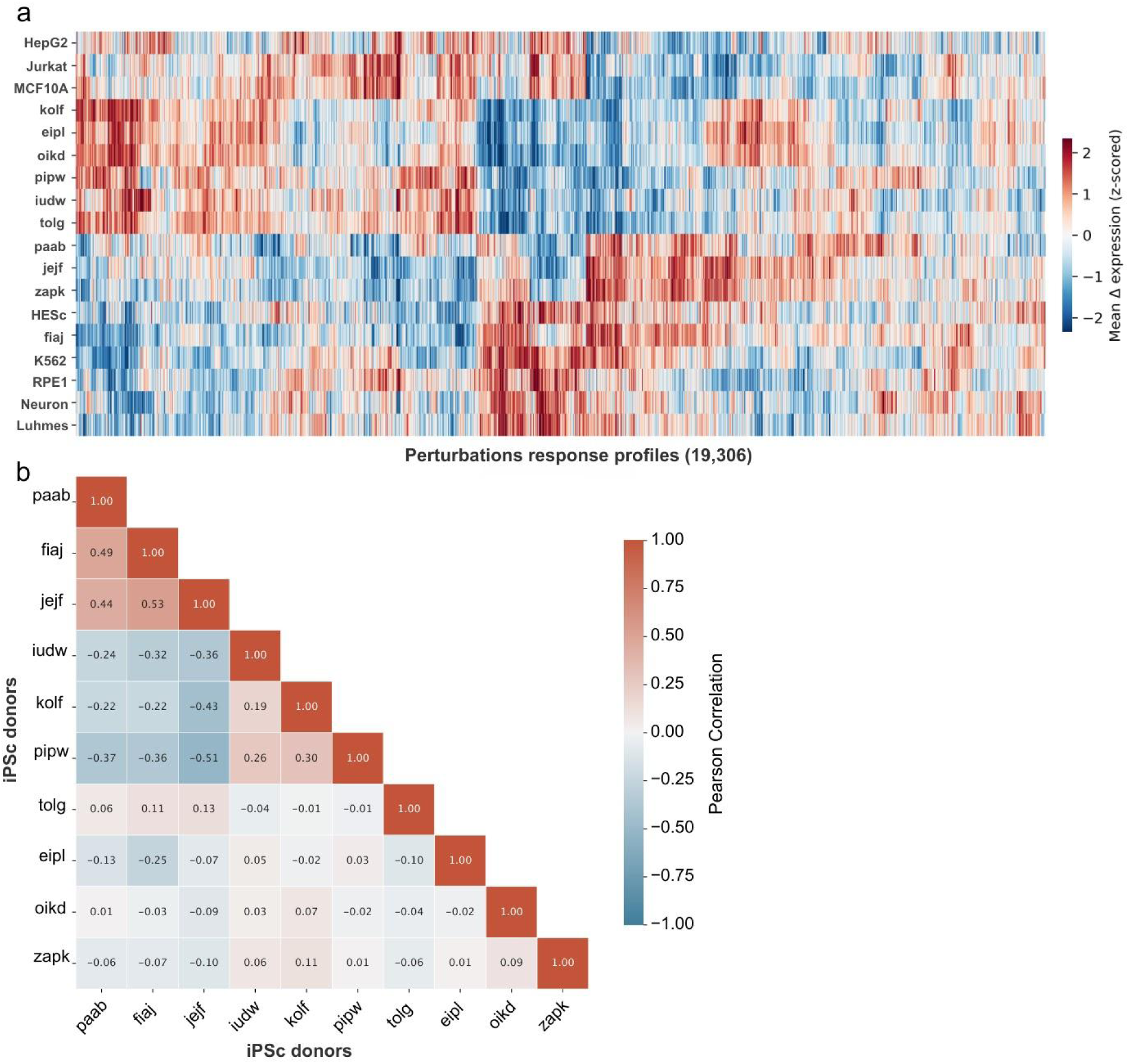
Global structure of the HyperMapDB perturbation response atlas. **a)** Heatmap showing mean perturbation response profiles across 18 cell types (rows) and 19,036 perturbations (columns). Colour scale indicates z-scored mean perturbation response, with red indicating upregulation and blue indicating downregulation. **b)** Pairwise Pearson correlation matrix of observed mean perturbation response profiles across the ten iPSC donors. Each cell shows the Pearson correlation coefficient between a pair of donors, with colour indicating the direction and magnitude of correlation. Positive correlations are shown in red and negative correlations in blue.

## Methods

### Overview of HyperMap

HyperMap considers a perturbation dataset of *N* cells, where *x_i_* ∈ *R^K^* is the baseline (control) gene expression vector for cell *i* with *K* genes, and *p_i_* denotes the genetic or chemical perturbation applied to that cell. Unperturbed cells have *p_i_* = ctrl. Each perturbation is represented either by a pretrained large language model (LLM) embedding, GenePT^29^(for gene perturbations), or a one-hot encoding (for drugs). The goal is to learn a function:

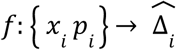

that maps the perturbation and its cellular context to a predicted transcriptional response 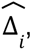, defined as:

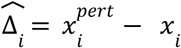

HyperMap is trained across multiple cellular contexts (cell types or donors), with each context treated as a distinct meta-learning task. This design enables rapid adaptation to a new cellular context using only a small number of observed perturbations.

### Data preprocessing and perturbation encoding

For each cellular context, a baseline control expression vector is computed as the mean expression across all control cells within that context. Perturbed expression vectors are normalised by subtracting this baseline to obtain context-specific Δ responses. Perturbation embeddings (GenePT or one-hot) are assigned to cells based on their condition labels. During batching, each perturbed cell is randomly paired with a control cell sampled with replacement from the same context to form matched training pairs.

### Cell and Perturbation Encoders

For each cell, a feed-forward encoder

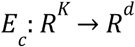

maps the control expression vector *x_i_* into a latent cell representation *z_c_*. A second encoder

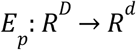

maps the perturbation embedding into a latent perturbation representation *z_p_*, where D is the dimensionality of the perturbation embedding. Both encoders consist of two hidden layers with LeakyReLU^50^ activations and dropout, projecting into a shared latent space of dimension *d* = 128.

### Context integration and dynamic parameter generation

HyperMap computes a context representation by attending the cell latent *z_c_* perturbation latent *z_p_*,

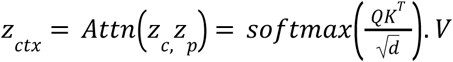

which captures how the perturbation interacts with the regulatory state of the cell. This context representation *z_ctx_* is then concatenated with *z_c_* and passed through a hypernetwork^28^ *H*_θ_,

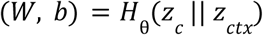

where *H* _θ_ is a two-layer feed-forward network that generates dynamic weights W and biases b for each specific cell-perturbation pair.

### Perturbation-response predictor

The predicted transcriptional change is obtained by applying the dynamically generated weights W and biases b to the perturbation latent *z_p_*,

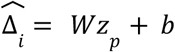

followed by a LeakyReLU non-linearity and a static linear projection into the full gene space of dimension *K*.

### Meta-learning strategy

Each reference cellular context is treated as a separate task. Training follows a MAML-style meta-learning framework^20^ implemented using the Higher library^51^, where at each outer iteration tasks are randomly sampled (n=4, with replacement if total tasks are fewer than 4). For each task, the model undergoes inner-loop adaptation across five batches, with five gradient update steps per batch. Meta-gradients are aggregated across the sampled tasks and used to update the shared parameters in the outer loop. During evaluation, the held-out new cellular context is adapted using 10–20 randomly selected seed perturbations, and the adapted model is then tested on the remaining unseen perturbations within that cell type.

### Loss function

Model parameters are optimised using mean squared error:

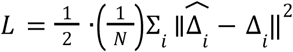

This error function trains the model to learn perturbation-specific transcriptional responses while controlling for baseline state via normalisation.

### Training setup

HyperMap was implemented in PyTorch^52^ and trained using Adam^53^ for the meta-optimiser (lr = 5×10⁻⁴) and SGD^54^. for inner-loop adaptation (lr = 1×10⁻²). Models were trained for 50 meta-epochs with a batch size of 512 and a meta-batch size of 4 tasks per outer update. Differentiable optimisation was performed using the Higher API^51^. In total, HyperMap comprises 6.6 million trainable parameters.

### Leave-one-out evaluation strategy

For all evaluations, HyperMap was assessed using a leave-one-out strategy, where the model is trained on all available contexts except one. The held-out context is then adapted using a small number of seed perturbations and evaluated on the remaining unseen perturbations within that context. Dataset-specific details, including the number of seed perturbations and evaluation perturbations, are described in the corresponding sections.

### Evaluation metrics

All metrics were computed by comparing predicted versus observed Δ expression for each perturbation. The top 20 genes per perturbation per donor were identified using the Wilcoxon rank-sum test comparing perturbed versus control cells, and ranked by absolute test statistic to capture both up- and down-regulated genes. Performance was quantified as the Pearson correlation between predicted and observed Δ expression restricted to these top-20 genes (top-20 Pearson). Gene identification accuracy was assessed using recall of top response genes (n=20), defined as the fraction of true top genes that appeared in the model’s predicted top response genes. Global reconstruction accuracy was quantified using mean squared error (MSE) and mean absolute error (MAE) over the top-20 Δ genes. Statistical significance was assessed using a two-sided Mann-Whitney U-test across perturbations (P < 0.05).

### iPSC perturb-seq dataset and preprocessing

We used the targeted perturb-seq dataset profiling iPSCs from ten human donors^30^, comprising 436 genetic perturbations across the ten backgrounds. Raw Unique Molecular Identifier (UMI) count matrices were processed to retain only cells with confidently assigned guides, ≥100 detected genes, and genes present in ≥3 cells, normalised to median total counts per cell, and log-transformed. Gene features were selected as the top highly variable genes across all donors, yielding a final feature space of 2,500 genes.

### iPSC control and perturbation similarity score calculation

Pairwise control state similarity between donors was computed by pooling observed unperturbed cells across all donors and calculating energy distances between donor groups using the scPerturb package^36^. Raw energy distances (d) were converted to similarity scores (s) using a Gaussian kernel *s* = exp(− *d*^2^ /(2σ^2^)), where σ is the standard deviation of the distance distribution, such that a score of 1 indicates identical transcriptional distributions and a score of 0 indicates maximally divergent distributions. Perturbation response similarity was computed as the Pearson correlation (r) of observed mean Δ expression vectors averaged across all shared perturbations, rescaled to [0, 1] as (r + 1) / 2, where 1 indicates identical perturbation responses across donors and 0 indicates maximally opposing responses. Both metrics were thus on a comparable [0, 1] scale to enable direct comparison.

### iPSC training strategy with HyperMap

HyperMap was trained and evaluated on the iPSC dataset following the leave-one-out strategy described above, with ten seed perturbations used for adaptation and the remaining 427 perturbations used for evaluation.

### Gene ontology analysis

Donors with a median perturbation prediction performance below 0.5 (Pearson-DE20) were selected for GO enrichment analysis. For these donors, perturbations were ranked by their median Pearson-DE20 performance across the selected donors, and strongly versus weakly transferable genes were defined as the top and bottom quartiles, respectively. GO Biological Process enrichment was performed separately for each group using Enrichr^55^ (GO Biological Process 2023, Human^56^) via the GSEApy package^57^, and terms with FDR < 0.001 in either group were retained for visualisation.

### SciPlex dataset processing and testing strategy

The SciPlex3 dataset, comprising 188 small molecule compounds profiled across three tumour cell lines (A549, lung epithelial; K562, hematopoietic; MCF-7, breast epithelial), was used for drug transfer^6^. Each dataset was independently filtered to retain cells with ≥100 detected genes and genes present in ≥3 cells, normalised to median total counts, and log-transformed. Only cells treated at 0 μM (vehicle control) or 10 μM (treatment) doses were retained. Gene features were selected by taking the union of the top highly variable genes per cell line, combined with the top most highly expressed genes across the pooled dataset, yielding a final feature space of 2,587 genes. Drug perturbations were represented as one-hot encodings. HyperMap was trained and evaluated following the leave-one-out strategy described above, with 20 randomly selected seed perturbations used for adaptation and the remaining 168 drug perturbations used for evaluation.

### Genetic cell-line dataset and training strategy

Four human cell lines spanning distinct lineages were used: RPE1 (retinal epithelial), K562 (hematopoietic), HepG2 (hepatic) and Jurkat (lymphoid), sourced from two independent perturb-seq studies^7,35^. Each dataset was independently filtered to retain cells with ≥100 detected genes and genes present in ≥3 cells, normalised to median total counts, and log-transformed. Gene features were selected by taking the union of the highly variable genes from each cell line, combined with the top-most highly expressed genes across the pooled dataset, yielding a final feature space of 2,947 genes. Only perturbations shared across all four cell lines were retained for cross-cell-line evaluation (n=1,742). HyperMap was trained and evaluated following the leave-one-out strategy described above, with 20 randomly selected seed perturbations used for adaptation and the remaining 1,722 perturbations used for evaluation.

### GEARS, scPRAM and scGPT training setup

GEARS is a graph-based deep learning model that integrates a Gene Ontology knowledge graph with gene embeddings to predict transcriptional responses to genetic perturbations^9^. scPRAM is a VAE-based model that uses optimal transport to match control and perturbed cells in latent space and an attention mechanism to compute perturbation vectors for held-out cells^31^. scGPT is a transformer-based foundation model pretrained on 33 million human single cells, fine-tuned for perturbation prediction^15^. All three models were trained using the default recommended parameters. As GEARS and scGPT lack a cross-context transfer mechanism, they were provided with the same number of perturbations in the held-out context as HyperMap’s seed perturbations. scPRAM supports cross-context transfer and was therefore trained on the full reference context data and transferred to the held-out context, matching HyperMap’s training strategy. All models were evaluated on the same held-out perturbations following the leave-one-out strategy described above.

### Number of seed perturbations and seed perturbation type

To assess the effect of seed perturbation number on adaptation performance, HyperMap was evaluated across a range of seed set sizes (1–100 perturbations) for each held-out donor, with performance averaged over 10 independent runs with randomly sampled seed perturbations. To assess the effect of seed perturbation type, four selection strategies were compared against a random baseline: least consistent (highest coefficient of variation across training donors), most consistent (lowest coefficient of variation among high-effect perturbations), and pathway-specific (iPSC-relevant transcription factors and chromatin regulators: POU5F1, PRDM14, ZNF281, STAT3, EZH2, SETD1A, CHD4, BCOR, ARID1A, DGCR8). Performance for each strategy was evaluated using 10 seed perturbations and reported as percentage change relative to random selection, averaged across donors, using the evaluation metrics described above.

### Training time

Model training time was benchmarked as a function of the number of perturbations (n = 100, 300, 700, 1,500) using synthetic datasets with a fixed number of cellular contexts (n = 10), 2,500 genes, and 10,000 observations per context (10% control cells). Measured time included both model-internal data preprocessing and training. Each model was benchmarked on a single NVIDIA Tesla V100-SXM2 (32GB) GPU under identical conditions. HyperMap, GEARS, scPRAM and scGPT were each benchmarked independently and training time was compared across models at each perturbation count.

### iPSC extension to unseen perturbations

To evaluate HyperMap’s ability to predict responses to perturbations not observed in any training context, the iPSC dataset was used in a modified leave-one-donor-out design in which HyperMap was trained on nine donors using a restricted set of 200 perturbations, with the remaining 236 perturbations withheld from all training donors. The held-out donor was adapted using 10 seed perturbations drawn only from the 200 training perturbations, and the model was evaluated on the 236 withheld perturbations and 18,609 protein-coding genes from GENCODE v47^58^ that overlapped with the GenePT embedding vocabulary, yielding a total of 19,036 predicted perturbation profiles per donor. GEARS and scGPT, which also support unseen perturbation prediction, were evaluated on the same 236 withheld perturbations under identical training restrictions for comparison.

### HyperMapDB data collection and merging

Perturb-seq datasets were curated from independent published screens spanning 18 cellular contexts: HepG2^35^, hESC^38^, Jurkat^35^, K562^7^, Luhmes^40^, MCF10A^41^, derived neurons^42^, RPE1^7^, and 10 iPSC donors^30^. Each dataset was independently filtered to retain cells with ≥100 detected genes, genes present in ≥3 cells, and single-gene perturbations only, excluding combinatorial perturbations. Data were normalised to median total counts and log-transformed. To harmonise gene features across all contexts, datasets were subset to the intersection of genes present in all 18 contexts. To balance cell numbers across contexts and perturbations, and consistent with HyperMap’s ability to learn effectively from sparse data, control cells were capped at 300 per context and perturbed cells were capped at 150 cells per perturbation per context. The merged dataset was then subset to the top highly variable genes across all contexts, yielding the final feature space of 2,500 genes.

### HyperMapDB data expansion

HyperMap was retrained on the full curated catalogue using all 18 cellular contexts as training tasks. For each context, the model was adapted using up to 20 randomly selected seed perturbations from the experimentally observed data, with the number of seeds capped by the available perturbations per context. Predictions were then generated for all 19,036 protein-coding genes from GENCODE v47^58^ overlapping the GenePT embedding vocabulary, yielding a complete 18 × 19,036 context-by-perturbation matrix (n = 342,648 context-by-perturbation combinations).

### HyperMapDB t-SNE visualisation

Pseudobulk Δ expression profiles were computed by averaging predictions per perturbation per context and z-scored within each context. The first principal component, dominated by knockdown of core ribosomal proteins (RPS/RPL family) and reflecting a global essentiality axis, was regressed out prior to all subsequent analyses, consistent with the known dominance of essential gene perturbations in large-scale perturb-seq screens^7^. Perturbation profiles summarised across all 18 cell types were then clustered using Leiden clustering (resolution=0.8) on principal components, with cluster labels assigned using GO Biological Process enrichment via Enrichr^55^. All 342,648 perturbation-cell type pairs were embedded using openTSNE^59^, with each point representing one perturbation in one cell type, colored by the perturbation cluster labels derived above.

### HyperMapDB perturbation effect size and cell-type specificity

For each of the 19,036 perturbations, two summary statistics were computed across all 18 cell types. Perturbation magnitude was defined as the mean absolute Δ expression across all cell types from the pseudobulk matrix. Cell-type specificity was quantified as the fraction of cross-cell-type variance remaining after removing the global essentiality axis (PC1), such that a low score indicates a consistent response across cell types and a high score indicates a predominantly cell-type-specific response. Perturbations were classified into four categories using median splits on both axes: strong and consistent, strong and variable, weak and consistent, and weak and variable.

## Data availability

The HyperMapDB resource, comprising transcriptomic perturbation response profiles for 19,036 gene knockdowns across 18 cell contexts, is available on Figshare at 10.6084/m9.figshare.31831081. All perturb-seq datasets used for model training and evaluation are publicly available: the iPSC perturb-seq^30^, the SciPlex3 dataset^6^, and the essential gene knockdown dataset^7,35^. Additional datasets used to construct HyperMapDB are listed in Extended Table 2 with their respective accession identifiers.

## Code availability

HyperMap code and example training scripts are available on GitHub: https://github.com/bhavya1929/HyperMap.

## Acknowledgements

This work was supported by Science Foundation Ireland [18/CRT/6214 to B.D.], through the Centre for Research Training in Genomics Data Science, the EU Horizon 2020 research and innovation programme under the Marie Skłodowska-Curie grant agreement [945385 to B.D.], and the Bridge2AI: Cell Maps for AI (CM4AI) Data Generation Project (NIH OT2 OD032742). The authors thank members of the Ideker Lab at UC San Diego and the GoldLab at University College Dublin for helpful discussions.

## Author Contributions

B.D., J.G., and T.I. designed the study and developed the conceptual ideas. B.D. collected all input datasets, performed data processing and model development. B.D. implemented the model and all computational analyses under supervision of J.G. and T.I. B.D., J.G., and T.I. wrote the manuscript

