## extended table for "HyperMap: An Efficient Framework for Transferring Perturbation Responses Across Diverse Biological Contexts"

| Table | Title |
|---|---|
| Extended Table 1 | GO Biological Process enrichment analysis for knockdowns that transfer well and do not transfer well across cellular contexts |
| Extended Table 2 | Studies and datasets included in HyperMapDB |

**Extended Table 1 |** GO Biological Process enrichment analysis for knockdowns (KD) which transfer well and do not transfer well

| GO Term | Category | Gene Overlap (KD transfers well) | Gene Set Size (KD transfers well) | Adjusted P-value (KD transfers well) | Gene Overlap (KD does not transfer well) | Gene Set Size (KD does not transfer well) | Adjusted P-value (KD does not transfer well) |
|---|---|---|---|---|---|---|---|
| RNA Processing | 0 | 14 | 183 | 1.21E-09 | 4 | 183 | 1.51E-01 |
| RNA Pol II Elongation Regulation | 0 | 8 | 77 | 2.83E-06 | 4 | 77 | 2.09E-02 |
| snRNA Processing | 0 | 5 | 19 | 7.16E-06 | 0 | 0 | 0.00E+00 |
| RNA 3' End Processing | 0 | 6 | 38 | 7.16E-06 | 1 | 38 | 2.93E-01 |
| Translation | 1 | 9 | 234 | 3.25E-04 | 12 | 234 | 2.29E-06 |
| Macromolecule Biosynthetic Proce | 1 | 8 | 183 | 3.94E-04 | 11 | 183 | 2.29E-06 |
| Gene expression | 1 | 13 | 296 | 2.83E-06 | 11 | 296 | 4.79E-05 |
| mRNA Splicing, via spliceosome | 1 | 11 | 211 | 4.58E-06 | 10 | 211 | 2.11E-05 |
| mRNA Processing | 1 | 11 | 214 | 4.58E-06 | 9 | 214 | 1.32E-04 |
| Transcription Initiation by RNA Pol | 2 | 2 | 52 | 1.98E-01 | 7 | 52 | 3.43E-06 |
| DNA Transcription Initiation | 2 | 2 | 57 | 2.01E-01 | 7 | 57 | 3.98E-06 |
| RNA Pol II Initiation Regulation | 2 | 2 | 56 | 1.98E-01 | 7 | 56 | 3.98E-06 |

**Extended Table 2 |** Studies and datasets included in HyperMapDB

| Cell Type / Context | Study | Perturbations after filtering |
|---|---|---|
| HepG2 | Nadig et al. 2025 | 2317 |
| Jurkat | Nadig et al. 2025 | 2317 |
| K562 | Replogle et al. 2022 | 1804 |
| RPE1 | Replogle et al. 2022 | 2013 |
| hESC | Genga et al. 2019 | 50 |
| LUHMES | Lalli et al. 2020 | 14 |
| MCF10A | Hill et al. 2018 | 27 |
| Neurons | Tian et al. 2019 | 24 |
| iPSC (eipl) | Nourreddine et al. 2024 | 436 |
| iPSC (fiaj) | Nourreddine et al. 2024 | 436 |
| iPSC (iudw) | Nourreddine et al. 2024 | 436 |
| iPSC (jejf) | Nourreddine et al. 2024 | 436 |
| iPSC (kolf) | Nourreddine et al. 2024 | 436 |
| iPSC (oikd) | Nourreddine et al. 2024 | 436 |
| iPSC (paab) | Nourreddine et al. 2024 | 436 |
| iPSC (pipw) | Nourreddine et al. 2024 | 436 |
| iPSC (tolg) | Nourreddine et al. 2024 | 436 |
| iPSC (zapk) | Nourreddine et al. 2024 | 436 |
